# The role of the *Trypanosoma cruzi* enzyme L-threonine 3-dehydrogenase in combating stressful environments

**DOI:** 10.1101/2025.09.08.674805

**Authors:** Ana María Mejía-Jaramillo, Paola García-Huertas, Carlos Renato Machado, Omar Triana-Chávez

## Abstract

*Trypanosoma cruzi* is a digenetic parasite that undergoes various transformations to complete its life cycle. Changes between hosts and vectors involve exposure to stressful environments, for which it has developed different strategies to cope with such stress. L-threonine 3-dehydrogenase (TDH) is a key enzyme in trypanosome metabolism, and several studies have shown that inhibiting TDH affects parasite survival. To understand the role of TDH in *T. cruzi*, we investigated its expression in different benznidazole (Bz)-resistant clones and overexpressed it in a Bz-susceptible clone. After overexpressing TDH and exposing it to reactive oxygen species (ROS), alkylating agents, and drugs such as Bz, we evaluated certain biological parameters. Our results show that TDH led to higher survival rates when exposed to H_2_O_2_ and increased tolerance to Bz. Moreover, the parasites were able to infect more cells, and their mitochondrial membrane potential (Ψm) remained unchanged, both of which are linked to higher tolerance to ROS. Finally, parasites overexpressing TDH were less vulnerable to genetic damage caused by agents such as MMS and gamma radiation. Overall, our results demonstrate that TDH, a key enzyme in threonine metabolism, helps combat stressful environments and, under certain experimental conditions, supports the survival of parasites.

## Introduction

L-threonine 3-dehydrogenase (TDH) is an enzyme found in various bacteria, fungi, protozoa, and some animals. In humans, its activity is very low o absent in most tissues [1,2]. TDH is primarily located in the mitochondria and plays a role in the catabolism of the amino acid L-threonine, converting it to 2-amino-3-ketobutyrate, which is then typically transformed into glycine and acetyl-CoA utilizing NAD+ as a cofactor and producing NADH. The formed 2-amino-3-ketobutyrate can spontaneously decarboxylate or be converted by aminoacetone synthase into aminoacetone, which can then enter additional metabolic pathways, such as pyruvate metabolism or the methylglyoxal pathway [3,4]. Methylglyoxal is a highly reactive dicarbonyl that contributes to lipid peroxidation, protein glycation, and DNA damage, leading to oxidative and carbonyl stress in cells. Furthermore, elevated NADH levels can lead to an over-reduction of the electron transport chain, resulting in the leakage of electrons and the generation of reactive oxygen species (ROS), particularly superoxide [4].

It was recently demonstrated that inhibiting TDH in *Trypanosoma cruzi* reduces glycine and acetate production, thereby disrupting parasite growth and viability, suggesting that TDH may be crucial for the survival of specific pathogens [1]. Additionally, Ochoa-Martínez et al. (2025) analyzed the expression of 13 genes by digital PCR in four Mexican *T. cruzi* isolates treated with benznidazole (Bz) and nifurtimox (Nfx) and found that TDH was notably overexpressed in these parasites [5]. Moreover, using proteomic methods, Andrade et al. (2008) found that TDH was exclusively expressed in induced *in vivo* benznidazole-resistant parasites [6].

It is well known that under drug-induced stress (e.g., exposure to antibiotics, antifungals, or chemotherapeutic agents), pathogens frequently experience oxidative and metabolic stress, as well as the disruption of key biosynthetic pathways [7]. In these situations, TDH may become essential as it helps the pathogen redirect metabolism to sustain energy production and biosynthesis when other pathways are blocked. Moreover, TDH is vital for maintaining redox homeostasis due to the reaction it catalyzes, which involves NAD⁺/NADH and contributes to the redox balance, often disrupted by drug stress [1]. Some pathogens upregulate TDH in response to drug pressure as part of a broader stress response network [5,6]. In this sense, we hypothesized that TDH could be a key enzyme in response to Bz and oxidative stress in *T. cruzi*. To address these questions, we examined various features of TDH in trypanosomes and its potential impact on the response to oxidative stress and genetic damage. We overexpressed TDH in a Bz-susceptible *T. cruzi* clone and evaluated the response to alkylating agents, reactive oxygen species (ROS), and drugs such as Bz. Additionally, we knocked down TDH in *T. brucei* and determined the localization of the protein in this parasite. Our results provide new insights that will help elucidate the mechanisms underlying the *T. cruzi* response to metabolic stress induced by various external stimuli.

## Materials and Methods

### Parasites

The *Trypanosoma cruzi* TcI 61S clone 11, which is susceptible to Bz with an Inhibitory Concentration 50 (IC_50_) of 11.7 µM, was used in this study [8,9]. Additionally, a Bz- and Nfx-resistant clone (61R clone 3) obtained by Mejia et al. (2012) with an IC_50_ of 54 µM was also used [8]. Epimastigotes were cultivated in supplemented Roswell Park Memorial Institute (RPMI) 1640 medium or in Liver Infusion Tryptose (LIT) medium supplemented with 10% (vol/vol) heat-inactivated Fetal Bovine Serum (FBS) at 28 °C. Cultures were maintained in exponential growth by passage every seven days. Transfected *T. cruzi* parasites were maintained with 100-250 μg/ml of G418. *T. brucei* BSF (Lister 427; clone 221a) were grown in HMI-9 medium [10] at 37 °C under a 5% CO_2_. *T. brucei* BSF lines (2T1) that constitutively express T7 RNA polymerase and the tetracycline repressor protein [11] were grown in medium containing 2 μg/ml phleomycin. Transformed *T. brucei* were maintained in medium with 5 μg/ml blasticidin or 2.5 μg/ml hygromycin.

### Construction of trypanosomal vectors and transfections

All primers used in this work are described in Table S1. The full-length ORF of L-threonine 3-dehydrogenase (BCY84_00949, TcTDH) from *T. cruzi* was cloned into the pTEX [12] and pTREX [13] vectors. The vectors used for localization and knock-down experiments in *T. brucei* were generated as reported in [14]. The 999 bp fragment (Tb927.6.2790), lacking the stop codon, was digested with *Hind*III/*Xba*I and ligated into the pIS^BLA^RPI^12myc^ vector [11] in frame with 12 copies of the c-Myc epitope. Ligation was performed such that an epitope (9E10) derived from the human c-myc protein was added to the C terminus of TbTDH. After cleaving the *Pst*I site within the TbTDH fragment, the construct was used to transfect BSF *T. brucei* to tag the endogenous TbTDH allele. For the TDH knock-down experiments, copies of the full-length *T. brucei* gene were cloned in opposite orientations into the pRPa^iSL^ vector. The *Asc*I digested construct was electroporated into *T. brucei* 2T1 parasites [11]. All the plasmids were confirmed by sequencing.

### Parasite transfections

Electroporation of *T. cruzi* epimastigotes and *T. brucei* BSF (WT and 2T1 lines) were done using the Amaxa® Cell Line Nucleofector® Kit T from Lonza (Catalog number VCA-1002, Lonza, Basel, SW) with the X-001 program in the Amaxa nucleofector II b (Lonza, Basel, SW) [15,16]. *T. cruzi* was selected with G418 (100 μg/ml), and pTREX-transfected parasites were seeded in 96 well-plates. Parasites transfected with pTEX-GFP or empty pTREX served as controls. *T. brucei* transfectants were selected with blasticidin or hygromycin. Overexpression in *T. cruzi* was verified by RT-qPCR [9], Northern blot, or western blot [15,16]. Knockdown of TDH in *T. brucei* was verified by Northern blot.

### Determination of the median inhibitory concentration (IC_50_) for benznidazole

Exponential phase *T. cruzi* epimastigotes were prepared at a final concentration of 5 × 10^5^ parasites/ml in medium with 10% FBS. In each assay, 10 concentrations of the drugs were evaluated, and each concentration was assessed four times in 96-well plates. Untreated parasites were used as growth controls. After incubation for 72 h, 20 μl of alamarBlue was added to each well, and plates were incubated for an additional 16 h. The cell density of each culture was determined as described previously [16], and the IC_50_ was established. One-way ANOVA with Tukey’s multiple comparison tests was used to determine differences between IC_50_. Values were expressed as mean ± SD, and statistical significance was determined at p<0.05.

### Survival curves

1 × 10^6^ or 1 × 10^7^ epimastigotes/ml in logarithmic phase were seeded to monitor cumulative parasite growth over several days. To test benznidazole sensitivity, parasites were treated with 60, 120, and 240 µM benznidazole and followed for seven days. The hydrogen peroxide (H_2_O_2_) response was assessed by treating parasites with 50, 100, and 150 µM H_2_O_2_ for 20 minutes, followed by resuspension in fresh medium and counting after 72 hours. To test the growth of parasites treated with the DNA alkylating agent methyl methanesulfonate (MMS), parasites were exposed to 1.5 mM MMS for one hour, resuspended in fresh medium, and counted daily for six days. For the gamma-irradiated parasite growth curve, cells were exposed to a dose of 500 Gy (1578 Gy/h for 20 minutes) in a cobalt-60 (60Co) irradiator located at the Centro de Desenvolvimento da Tecnología Nuclear (CDTN), Belo Horizonte, Brazil. After irradiation, cells were incubated and counted daily for 13 days. In all experiments, cell numbers were determined in a cytometry chamber using the erythrosine vital stain (PBS 1x + erythrosine 0.4%) to differentiate live from dead cells.

To test the essentiality of the TDH gene, *T. brucei* parasites were seeded at 1 × 10 ^ ^5^ cells/mL and incubated at 37 °C in the presence or absence of tetracycline (1 μg/mL). Every 24 hours, parasite growth was monitored microscopically, and the culture was diluted back to 1 × 10^5^ cells/mL [17]. Experiments were performed in triplicate, and non-treated controls were used in all cases. The differences between treated and untreated parasites were determined using different statistical tests in GraphPad Prism 10 software.

### Flow cytometry analyses

1 × 10^7^ parasites/ml were treated with 120 μM Bz or non-treated and incubated at 28 °C for 48 hours. Cells were washed with phosphate-buffered saline (PBS) and fixed with 70% ethanol at 4 °C for 24 hours for the cell cycle analysis. Subsequently, cells were washed once and resuspended in PBS containing 10 μg/ml of propidium iodide and 10 μg/ml of RNAse A. After 45 min of incubation at 37 °C, the parasites were analyzed using BD FACSCalibur equipment. For mitochondrial membrane potential, parasites were washed and resuspended in PBS containing 0.8 nM 3,3’-dihexyloxacarbocyanine iodide (DiOC6). The suspension was incubated in the dark at room temperature for 15 minutes before cytometric analysis. Data analyses were made using Flowjo vX.0.7 software, comparing values from treated and non-treated parasites.

### Infection assay

To generate trypomastigotes, Vero cells cultured in DMEM with 10% FBS at 37 °C in 5% CO_2_ were infected with epimastigotes in the stationary phase [18]. Trypomastigotes emerged between days 7 and 10 and were used to infect 30,000 cells at a ratio of three trypomastigotes per mammalian cell. After incubation at 37 °C for 24 hours, extracellular parasites were removed by several washes. After 48 hours, the cells were stained with Giemsa, and 500 cells were counted per treatment. The numbers of amastigotes and the percentages of infection were obtained from five experiments; each performed in triplicate. Differences between pTEX-GFP and pTEX-TcTDH were examined using an unpaired t-test in the GraphPad Prism 10 software.

### Immunofluorescence microscopy

Trypanosomes in logarithmic growth phase expressing TbTDH were suspended at 1 × 10^6^ cells/ml in medium containing 100 nM MitoTracker Red (Molecular Probes) and incubated at 37 °C for 5 min. Cells were washed twice in PBS, fixed in 2% paraformaldehyde in PBS, and air-dried on glass slides as described in [19]. Cells were labeled with mouse anti-c-Myc (9E10) antibody (Santa Cruz Biotechnology) (diluted 1:100) for one hour and secondary antibody (AlexaFluor 488-conjugated goat anti-mouse antibodies; Molecular Probes) (diluted 1:400) for 30 min in PBS-S, with 2% horse serum. Parasite DNA was stained with 200 pM DAPI (Sigma) in 50% glycerol/PBS, and slides were viewed using a Zeiss LSM 510 confocal microscope.

## Results

### TDH is downregulated in Bz-resistant parasites induced *in vitro*

Since several studies have indicated that TcTDH is overexpressed in Bz/Nfx resistant parasites [5,6], we wanted to verify whether the transcription of this gene was altered in our *in vitro* induced resistant clone (61Rcl3) maintained with or without Bz at 54 µM. Relative quantification was performed by RT-qPCR using the susceptible clone as a control and normalizing mRNA levels to the expression of the hypoxanthine-guanine phosphoribosyltransferase (HGPRT) gene [9]. We observed that the TcTDH transcript was 2.2 times lower in the resistant clone compared to the susceptible clone. Exposure to Bz slightly increased the TDH expression in the resistant clone, although it was statistically significant (S1A Fig). Additionally, we detected that the growth of this resistant clone was significantly slower compared to the susceptible one (S1B Fig). Furthermore, Mejia et al. (2012) reported that this clone had a lower infection rate in Vero cells and fewer amastigotes per cell [8].

To evaluate TDH expression in other resistant parasites, we searched the raw RNA-seq data published by Mejía Jaramillo et al. (2025) and Lima et al. (2023). We assessed the mRNA levels of this gene in the resistant parasites [16,20]. After obtaining the reads from the nine copies of this gene (C4B63_18g2, C4B63_18g5, C4B63_18g8, C4B63_18g11, C4B63_42g114, C4B63_42g118, C4B63_42g121, C4B63_42g124, C4B63_42g127), we performed differential expression analysis [15] using the *T. cruzi* Dm28c reference genome (2018). For the first resistant clone (61R_cl4), there were no differences in expression (log2FC = −0.247, padj = 0.467), but for the second (LER), we observed a significant down-regulation (69%) compared to the control (log2FC = −1.675, padj = 1.0 × 10⁻²⁰) (S1 File). Finally, we searched for mutations in the sequences of the TcTDH gene in the susceptible and resistant parasites but did not encounter any in these clones (S1C Fig).

### Parasites overexpressing TDH were more susceptible to Bz

Given that our results of TDH gene expression in resistant clones contradicted previous reports, we overexpressed TDH in Bz-susceptible parasites to evaluate their response to Bz and other stressful stimuli. Overexpression of the pTEX-TcTDH-transfected parasites was confirmed by RT-qPCR (S2A Fig) and northern blot (S2B Fig), and by western blot (S2C Fig) in the pTREX-TcTDH parasites. Next, we determined the IC_50_ for Bz using alamarBlue after 72 hours of exposure to ten drug concentrations. The results showed that while parasites overexpressing TcTDH in the pTREX plasmid did not change their IC_50_ to Bz compared to the control, those transfected with the pTEX plasmid became more susceptible (S2D Fig). There were no differences in the growth curves between parasites transfected with pTEX and the susceptible clone used as a control (S2E Fig).

### pTEX-TcTDH parasites grow better with Bz

To evaluate the response to Bz more broadly, we exposed the pTEX-transfected parasites to different concentrations of Bz and quantified their growth daily. The pTEX-TcTDH parasites showed significantly higher growth than the control when exposed to 60 and 120 µM Bz on days 2 and 3. However, their growth began to decline and ultimately stopped on day 7. The pTEX-TcTDH parasites treated with 240 µM Bz began to die on day 3, and only a few cells were observed on day 7. However, pTEX-GFP parasites at the same concentration died completely after day 3, suggesting that TDH overexpression slightly increases tolerance to Bz (Fig 1A, S2F Fig).

**Fig 1.**
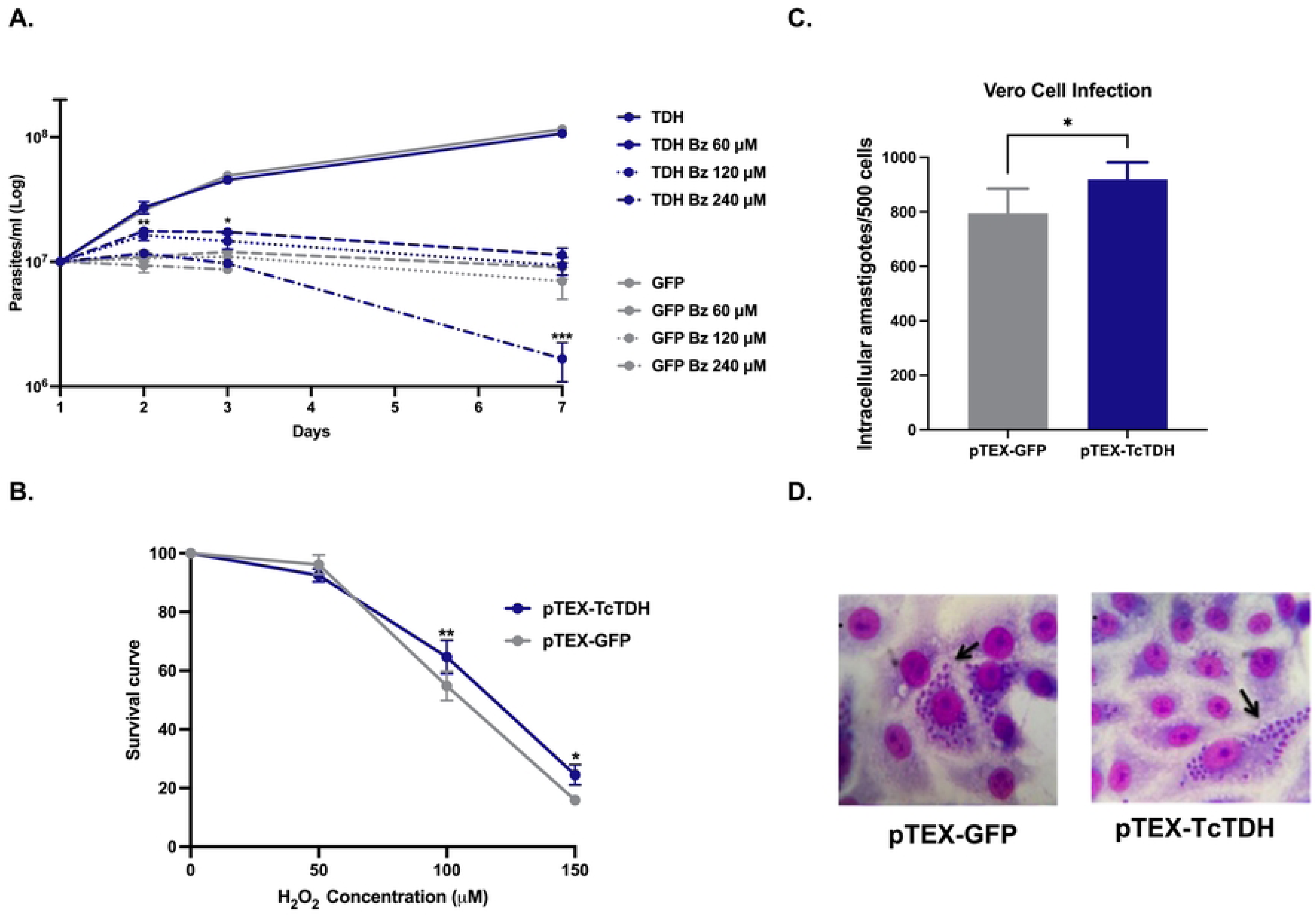
**A and B.** Survival curves of parasites overexpressing TDH compared to parasites transfected with GFP after treatment with 60, 120 and 240 µM of Bz for seven days (**A**) and treatment for 20 minutes with 50, 100 and 150 µM of H_2_O_2_ and quantification after 72 hours (**B**). **C and D.** Infection of Vero cells with pTEX-GFP and pTEX-TcTDH and amastigotes quantification after 48 hours (**C**) stained with Giemsa (**D**). Arrows indicate intracellular amastigotes. Statistical significance was determined using a two-way ANOVA (**A, B**), with Sidak’s multiple comparison correction and an unpaired t test (**C**), performed in GraphPad Prism10. ***p < 0.001; **p < 0.01; *p < 0.05.

### Parasites overexpressing TcTDH survive in the presence of H_2_O_2_

To better understand the results obtained with pTEX-TcTDH-transfected parasites, a second experiment was performed in which the parasites were exposed to 50, 100, and 150 µM H_2_O_2_. The results indicated that pTEX-TcTDH parasites had higher survival rates to H_2_O_2_ than those found for pTEX-GFP (Fig 1B, S2G Fig).

Given that transfected parasites demonstrated improved growth in the presence of Bz and H_2_O_2_, we evaluated the infectivity of these parasites in Vero cells, as infection is associated with oxidative stress. As expected, the pTEX-TcTDH parasites displayed a higher infection rate (49%) compared to the pTEX-GFP parasites (27.5%). Likewise, the number of amastigotes per cell in the pTEX-TcTDH parasites was significantly higher than in the pTEX-GFP parasites (Figs 1C and 1D).

### Mitochondrial membrane potential and cell cycle are not altered in TcTDH-overexpressing parasites after treatment with Bz

To determine changes in mitochondrial function, we measured the mitochondrial membrane potential (Ψm) of Bz-treated and untreated parasites. While the pTEX-GFP parasites treated with Bz exhibited hyperpolarization of the mitochondrial membrane, represented by a mean fluorescence intensity of 501 compared to 276 in untreated parasites, pTEX-TcTDH parasites showed no change in mitochondrial membrane potential after treatment with Bz, with mean fluorescence intensities of 283 and 336 for Bz-treated and untreated parasites, respectively (Fig 2A).

**Fig 2.**
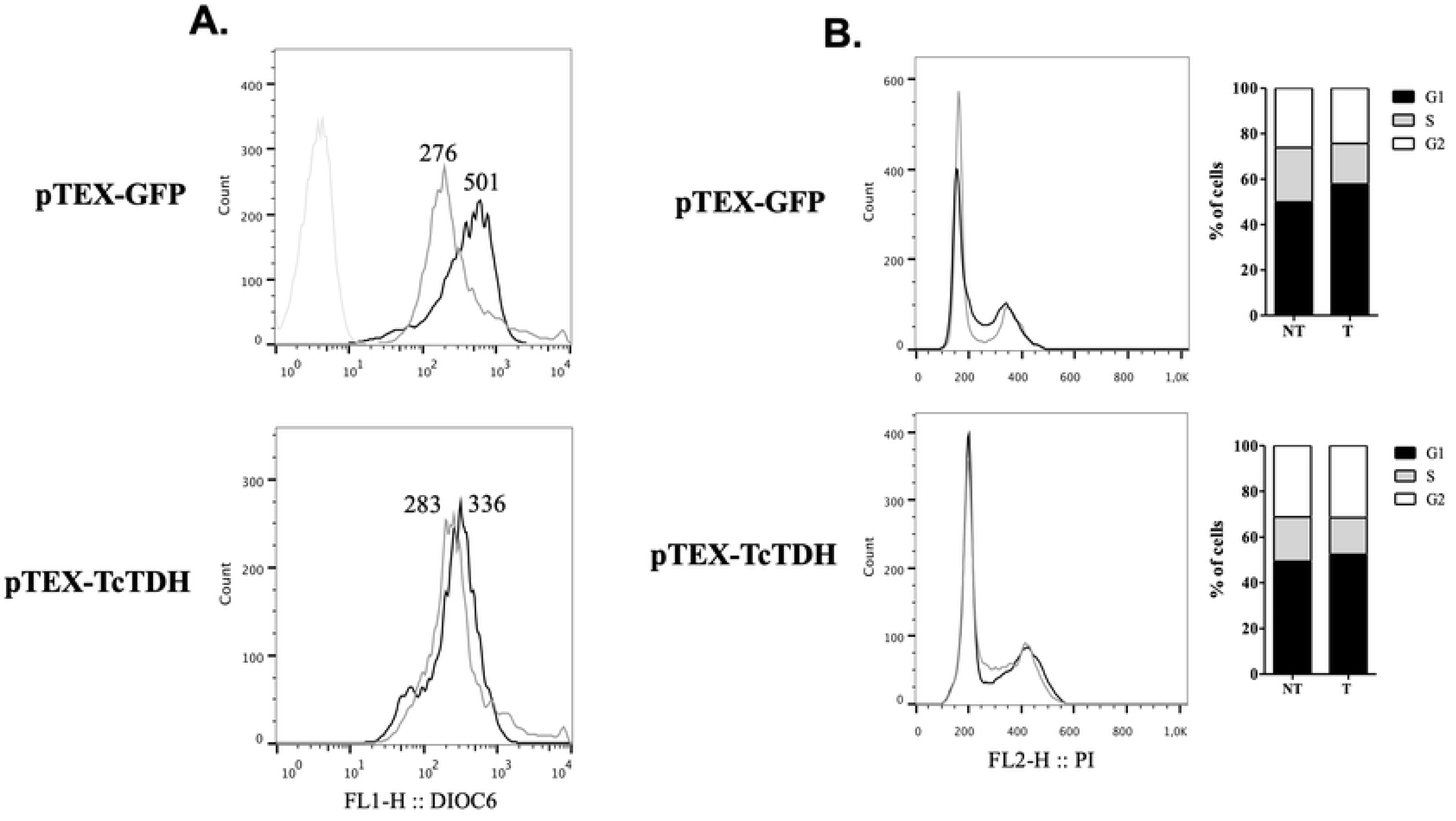
**A and B.** Flow cytometry analysis was performed to measure the mitochondrial membrane potential (ΔΨm) using DiOC6 staining (**A**), and to analyze cell cycle progression using propidium iodide (PI) (**B**), in parasites overexpressing either GFP or TcTDH. Parasites were either untreated (grey lines) or treated with 120 µM Bz (black lines). Readings were taken using a BD FACSCalibur Cell Analyzer flow cytometer.

Additionally, the effect of 120 μM Bz on the parasite cell cycle was assessed at 48 h post-treatment. pTEX-GFP parasites treated with Bz showed an accumulation of cells in the G1 phase, with about 8% more cells than untreated parasites, and a decrease in cells mainly in the S phase, where a difference of 5% was observed compared to non-treated parasites. pTEX-TcTDH parasites treated with Bz showed only 4% more cells in G1 than non-treated parasites and a difference of about 4% in the S phase (Fig 2B).

### Parasites overexpressing TcTDH are less susceptible to genetic damage

Genetic damage was analyzed to identify the response of pTEX-TcTDH parasites to agents such as MMS and gamma radiation. Following treatment with MMS (dotted lines), pTEX-TcTDH parasites exhibited greater growth than pTEX-GFP parasites from days 2 to 4 (Fig 3A). Additionally, the results obtained after irradiation showed statistically significant differences in growth kinetics between pTEX-GFP and pTEX-TcTDH parasites. Irradiated pTEX-TcTDH parasites (dotted lines) grew on day 2 and maintained a higher number of parasites during the assay compared to pTEX-GFP. After eight days, both pTEX-TcTDH and pTEX-GFP parasites doubled their growth. However, pTEX-TcTDH parasites reached a significantly higher number of parasites by day 13 (Fig 3B).

**Fig 3.**
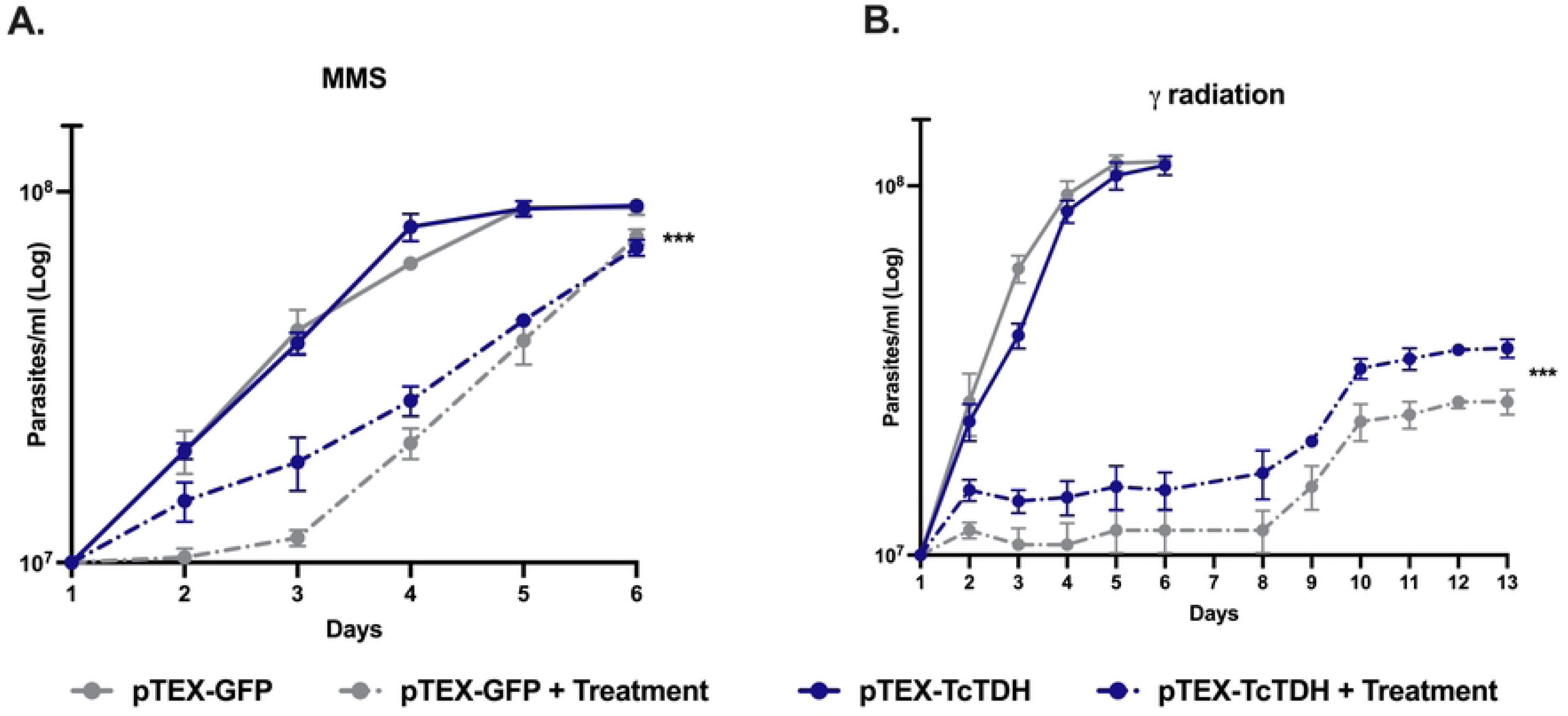
**A and B.** Survival curves of parasites overexpressing TDH compared to parasites transfected with GFP after treatment with 1.5 mM methyl methanesulfonate (MMS) for one hour and daily counts for six days (**A**) and after treatment with 500 Gy of gamma irradiation for 20 minutes and daily counts for 13 days (**B**). Statistical significance was determined using a two-way ANOVA performed in GraphPad Prism10. ***p < 0.001; **p < 0.01; *p < 0.05.

### TbTDH is not essential, and it is localized in the mitochondria

Considering that *T. cruzi* TDH is present in nine copies located in different alleles, gene deletion is a challenging process. Therefore, we decided to assess its essentiality in *T. brucei* using RNA interference (RNAi). We employed a tetracycline-inducible RNAi system to selectively knock down TbTDH transcripts, which was confirmed by northern blotting (S3A Fig). Growth experiments were conducted using two independent TbTDH RNAi lines. However, no significant differences in cumulative growth were observed after 12 days, regardless of the presence or absence of tetracycline, demonstrating that the TbTDH gene is not essential (S3B and S3C Figs). Finally, we implemented targeted integration to tag the endogenous TbTDH gene at the 3’ end with a sequence encoding the 9E10 epitope (derived from human c-Myc). The location of the carboxyl-tagged protein was determined using a monoclonal antibody and confocal microscopy, which revealed that the protein was confined to regions of the cell, as highlighted by the mitochondrion-specific dye Mitotracker (S3D Fig).

## Discussion

The L-threonine catabolism pathway is involved in different functions in both prokaryotes and eukaryotes, including energy production, homeostasis, and fatty acid synthesis [2]. In this study, we found that *T. cruzi* parasites overexpressing TDH, a key enzyme in threonine metabolism, are more tolerant to benznidazole, hydrogen peroxide, MMS, and gamma radiation. Likewise, we found that these parasites are more infectious and do not exhibit changes in their mitochondrial membrane potential; their progression through the cell cycle is also unaffected when exposed to Bz. Thus, our data suggest that the physiological role of L-threonine through the TDH pathway extends beyond the previously mentioned functions and sheds new light on the multiple tasks that this gene could have in the parasite that causes Chagas disease. For instance, the glycine produced from L-threonine via TDH is incorporated into trypanothione [21]. In this sense, understanding the role of trypanothione in drug metabolism, coupled with evidence of TDH overexpression in Bz-resistant parasites, led us to investigate the functional role of TDH in *T. cruzi*.

First, we examined TDH in other resistant phenotypes but found that it was either downregulated or unchanged. Although we observed that the addition of Bz to the culture increased TDH expression (S1A Fig), as previously reported [5], we decided to overexpress the enzyme in *T. cruzi* and determine the IC_50_ for Bz using alamarBlue since both results intrigued us. However, we obtained conflicting results. While overexpression with the episomal vector made the parasites more sensitive to Bz (S2D Fig), as seen with other resistant parasites expressing TDH at low levels, the integrative vector did not change the IC_50_ for Bz. The different outcomes may be due to a potential discrepancy in protein expression between the two vectors. However, a direct comparison of protein levels between parasites transfected with either vector was not possible because of the lack of suitable antibodies against TcTDH.

The previous finding indicated that the observed variations in TDH expression and Bz resistance might be attributable to enzyme expression levels, the drug concentrations used, or exposure time, among other factors. Thus, we selected the pTEX-TcTDH parasites to evaluate the response to Bz over several days, observing a significant survival of these parasites between days 2 and 3 compared to pTEX-GFP at 60 and 120 μM Bz. This demonstrates that TDH overexpression at certain Bz doses increases *T. cruzi* lifespan, at least in the first days. This period corresponded to the time during which Ochoa-Martínez et al. (2025) exposed parasites to Bz and subsequently quantified TDH expression [5].

In threonine catabolism, an imbalance in certain enzymes can result in the accumulation of 2-amino-3-ketobutyrate, a highly unstable product that spontaneously decarboxylates into aminoacetone [22]. Aminoacetone can undergo further oxidation by a class of enzymes called amine oxidases, which catalyze the oxidative deamination of the primary amine of the corresponding aldehydes, hydrogen peroxide, and ammonia. Aminoacetone undergoes oxidation to form MGO, which subsequently generates reactive oxygen species (ROS) in the form of hydrogen peroxide [4] (Fig 4).

**Fig 4.**
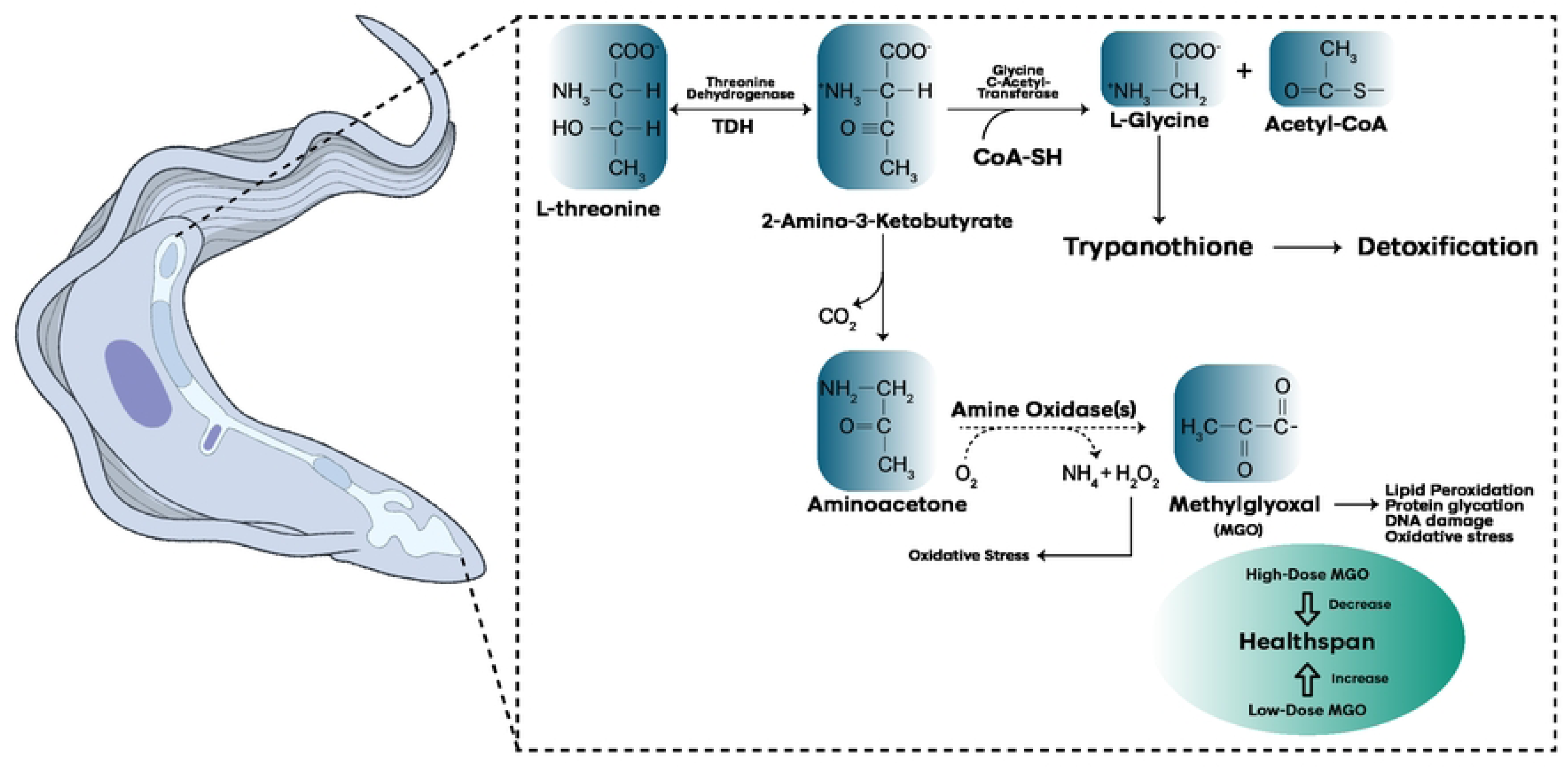
Summary of the TDH pathway and the most relevant toxic products. The level of expression of TDH and CoA-SH can favor either the detoxification pathway through trypanothione formation, or the generation of toxic products such as H_2_O_2_ and MGO. The concentrations of the latter, in turn, can favor an increase or decrease in the lifespan of trypanosomes.

The results obtained by Ravichandran et al. (2018) led us to hypothesize that the imbalance that we generated in this pathway by overexpressing TDH could generate greater adaptation to H_2_O_2_ and therefore cause the pTEX-TcTDH parasites to be more resistant to this compound [4]. Our findings support this hypothesis, as the parasites exhibited increased resistance to H_2_O_2_. In addition, the parasites were able to infect a greater number of cells, and their mitochondrial membrane potential (Ψm) remained unchanged, both of which are associated with higher tolerance to ROS. Grazielle-Silva et al. (2015) previously described this finding, determining that H_2_O_2_-resistant parasites were more infective than susceptible parasites [23].

Additionally, we wondered whether the parasites that were more tolerant to oxidative stress were also more resistant to alkylating agents and gamma radiation. Several studies have proposed that Bz acts by inducing oxidative stress and genetic damage. It has been proposed that the Bz induces double-stranded breaks in *T. cruzi* [5, 27]. Therefore, we exposed the pTEX-TcTDH parasites to MMS, an alkylating agent [28], and gamma radiation, which causes double-stranded DNA breaks [29]. We found that parasites overexpressing TcTDH were slightly resistant to genetic damage and showed no accumulation in any cell cycle phase. This indicates a relationship between resistance to genetic damage and TDH overexpression that could favor a response of DNA repair proteins and therefore result in tolerance to genetic damage and Bz [5]. This hypothesis must be addressed in future studies.

The observed variations in TDH expression and resistance to different external stimuli can be attributed to the findings reported by Ravichandran et al. (2018). The alteration of the threonine pathway in *C. elegans* due to the knock-out of Glycine C-Acetyl Transferase (GCAT) resulted in increased nematodes survival rates. This effect was also observed in the presence of low concentrations of MGO. Despite the assertion that MGO exerts a contradictory effect on organisms, affecting survival due to alterations in oxidative phosphorylation, ROS production, and changes in glycerol transport, among others. The authors hypothesized this phenomenon as a hormetic activity produced by MGO. Consequently, it has been observed that while low concentrations of this compound can result in increased survival, high concentrations can be lethal [4] (Fig 4). This same observation was described in *T. cruzi*, where Mesías et al. (2019) found that oxidative stress at high concentrations causes damage to biomolecules, but at low concentrations, oxidants are essential for cell signaling, and the oxidants/antioxidants balance may be able to trigger different cell fates [7].

The findings of this study suggest that this phenomenon may be a contributing factor of the observed variations in our results. If an imbalance is created in threonine pathway by mild overexpression of TDH, a slight accumulation of 2-amino-3-ketobutyrate may be generated, producing H_2_O_2_ and MGO in low concentrations. Consequently, parasites may exhibit enhanced response to the effects generated by Bz and increase their lifespan. However, it is plausible that TDH overexpression is associated with an increase in the GCAT enzyme, which would result in elevated levels of L-glycine, leading to a greater production of trypanothione, a compound that can detoxify and possibly enhance the parasites’ resistance to Bz (Fig 4). Future studies focused on the role of these compounds in Bz resistance are needed to address these questions.

Conversely, elevated TDH levels may result in a high accumulation of 2-amino-3-ketobutyrate and increased concentrations of H_2_O_2_ and MGO, leading to greater susceptibility to Bz, as was observed in the alamarBlue results for pTEX-TcTDH parasites. This could also explain why, in some resistant parasites, TDH levels are decreased. This reduction in enzyme levels could be a defensive strategy employed by the parasites to evade the generation of reactive species through the threonine catabolism pathway.

The results reported by Ravichandran et al. (2018) differed from those obtained in other studies, which they attributed to their specific experimental conditions [4]. The discrepancies regarding resistance to Bz and TDH levels in *T. cruzi* could also be explained by similar factors. Table 1 summarizes these features, showing that the strains used differed, as were the mechanisms of Bz resistance induction and IC_50_, the method used to evaluate the IC_50_, and the methodology used to quantify TDH levels. Furthermore, NTRI expression varies among strains, which has the potential to influence the way in which Bz is activated and the subsequent production of toxic compounds within the parasites.

**Table 1.** Summary of strains/clones and results obtained regarding resistance to Bz and TDH expression.

| Phenotype | Response to Bz | Bz IC<sub>50</sub> (μM) | TDH expression | FC | Methodology | Type of resistance | NTR expression | Reference |
| --- | --- | --- | --- | --- | --- | --- | --- | --- |
| CL-Brenner+Bz | Resistant | 56.5 | Up | 2.7 times | Digital PCR | Natural | Down | Ochoa-Martínez et al., 2025 |
| Sc4+Bz | Resistant | 76.0 | Same | N/A | Digital PCR | Natural | Up |  |
| CBu1+Bz | Partially resistant | 30.9 | Up | 15 times | Digital PCR | Natural | Same |  |
| Sum3+Bz | Partially resistant | 19.7 | Up | 1.04 times | Digital PCR | Natural | Down |  |
| LJ01+Bz | Susceptible | 9.6 | Up | 2 times | Digital PCR | N/A | Up |  |
| Clone27R | Resistant | N/D | Up | Exclusive expressed | Proteomic | *In vivo* induced | Same | Andrade et al., 2008 |
| 61R_cl3 | Resistant | 54 | Down | 2.2 times | qPCR | *In vitro* induced | Nonfunctional | Mejía et al., 2012 |
| 61R_cl3+Bz | Resistant | 54 | Down 61S/Up 61_cl3 | 2/1.1 times | qPCR | *In vitro* induced | Nonfunctional |  |
| 61R_cl4 | Resistant | 100.1 | Same | N/A | RNA seq | *In vitro* induced | Functional | Mejía-Jaramillo et al., 2025 |
| LER | Resistant | 220 | Down | 69% | RNA seq | *In vitro* induced | Down | Lima et al., 2023 |
| pTREX-TcTDH | Susceptible | 9.9 | Up | N/D | Western blot | N/A | Functional | This paper |
| pTEX-TcTDH | Susceptible | 5.5 | Up | 2.9 times | qPCR and NB | N/A | Functional |  |

N/A: not applicable; FC: Fold Change of TDH; Methodology: Technique used to quantify TDH expression.

Finally, although it has been suggested that TDH could be a good therapeutic target, our knock-down results in *T. brucei* showed that it is not an essential enzyme, so its inhibition with drugs may not be as relevant. However, in our resistant clone 61Rcl3, a decrease in fitness was observed that could be partly attributed to the downregulation of TDH, which is consistent with what has been reported by other authors [1]. Thus, it is possible that the experiments should be repeated in *T. cruzi*, where several drugs have a negative effect on the pathway [24,25].

## Conclusions

This study showed that *T. cruzi* parasites overexpressing TDH are more resistant to benznidazole, hydrogen peroxide, MMS, and gamma radiation. Consistent with this, we found these parasites are more infectious and do not show changes in their mitochondrial membrane potential; their progression through the cell cycle also remains unaffected when exposed to Bz. Therefore, TDH expression levels in trypanosomes could serve as an important trigger to help the parasite cope with environmental stress, including drug exposure.

## Acknowledgments

We are grateful to Professor John Kelly for allowing us to conduct the *T. brucei* experiments in their laboratory at The London School of Hygiene & Tropical Medicine (LSHTM) and Hader Ospina for their assistance with the bioinformatic analyses. Finally, we would like to thank Professor Frank Ávila for reviewing the English in the manuscript.

## Supporting information

**S1 Fig.**
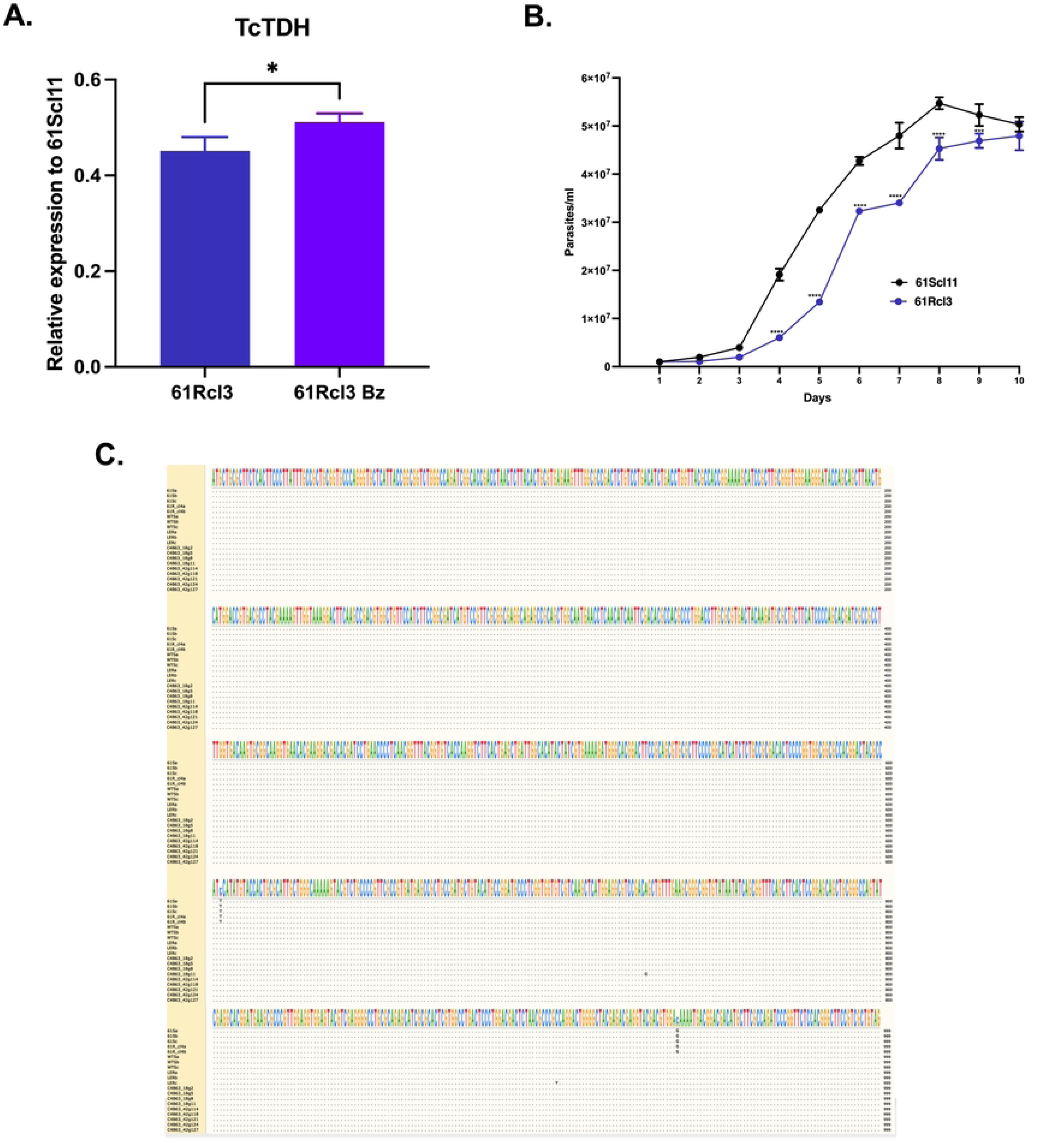
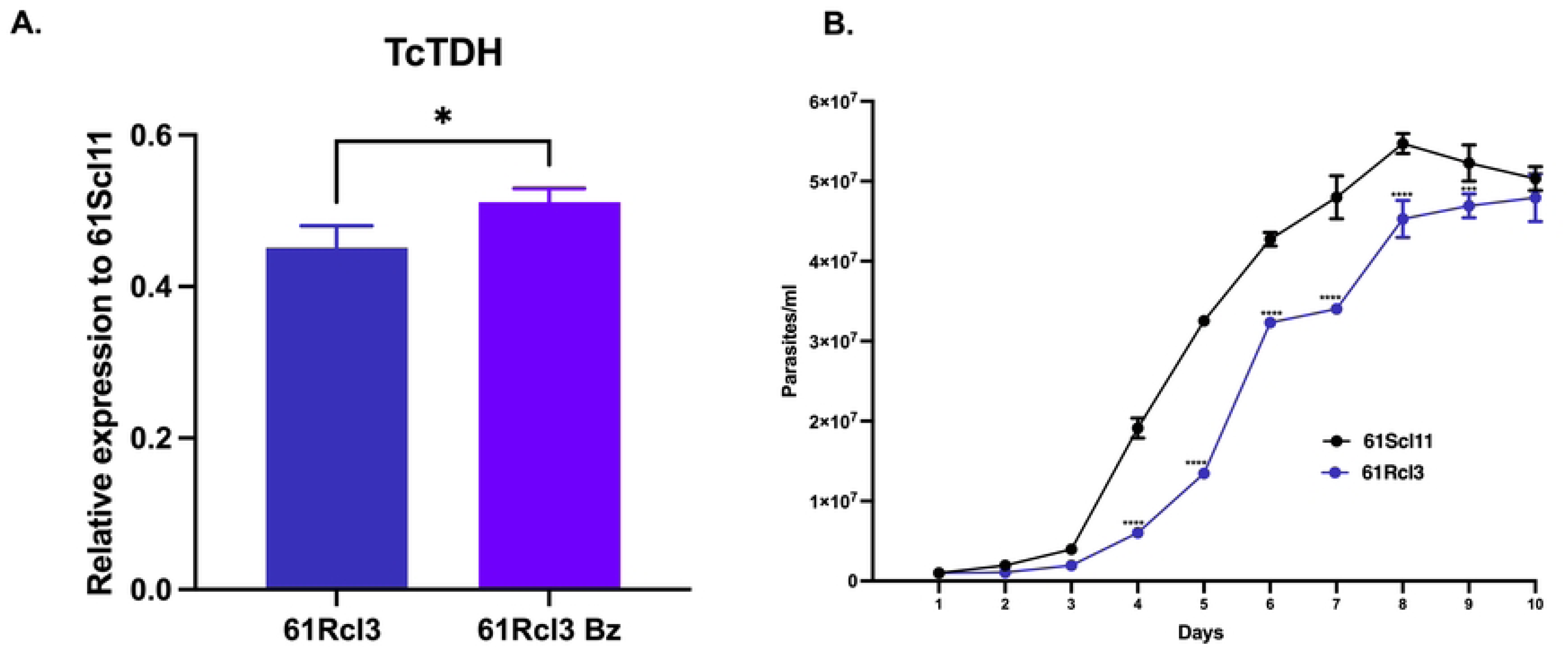
**A.** Relative quantification by RT-qPCR of TcTDH transcript in a resistant clone 61Rcl3 cultured without or in the presence of 54 µM of Bz (61Rcl3 Bz), compared with the sensitive clone (61Scl11). The mRNA levels were normalized using the expression of the reference gene (HGPRT). The relative quantification results were obtained using the REST2009 program by randomization test with 10,000 permutations (p < 0.05). The graph shows the data in triplicate and two independent experiments. **B.** The proliferation of epimastigotes was assessed by counting them in a Neubauer chamber every 24 hours for 10 days. Statistical significance was determined using a two-way ANOVA with Sidak’s multiple comparison test in GraphPad Prism 10. ****p < 0.0001; ***p < 0.001; *p < 0.05. **C.** Nucleotide alignment of the TcTDH gene from DNA sequencing of different *T. cruzi* clones. 61S and 61R_cl4: susceptible and resistant clones, respectively, taken from Mejía-Jaramillo et al., 2025. WTS and LER: susceptible and resistant clones, respectively, obtained from Lima et al., 2023 [16,20]. Copies of the TDH gene (C4B63_18g2, C4B63_18g5, C4B63_18g8, C4B63_18g11, C4B63_42g114, C4B63_42g118, C4B63_42g121, C4B63_42g124, C4B63_42g127) from *T. cruzi* Dm28c reference genome (2018).

**S2 Fig.**
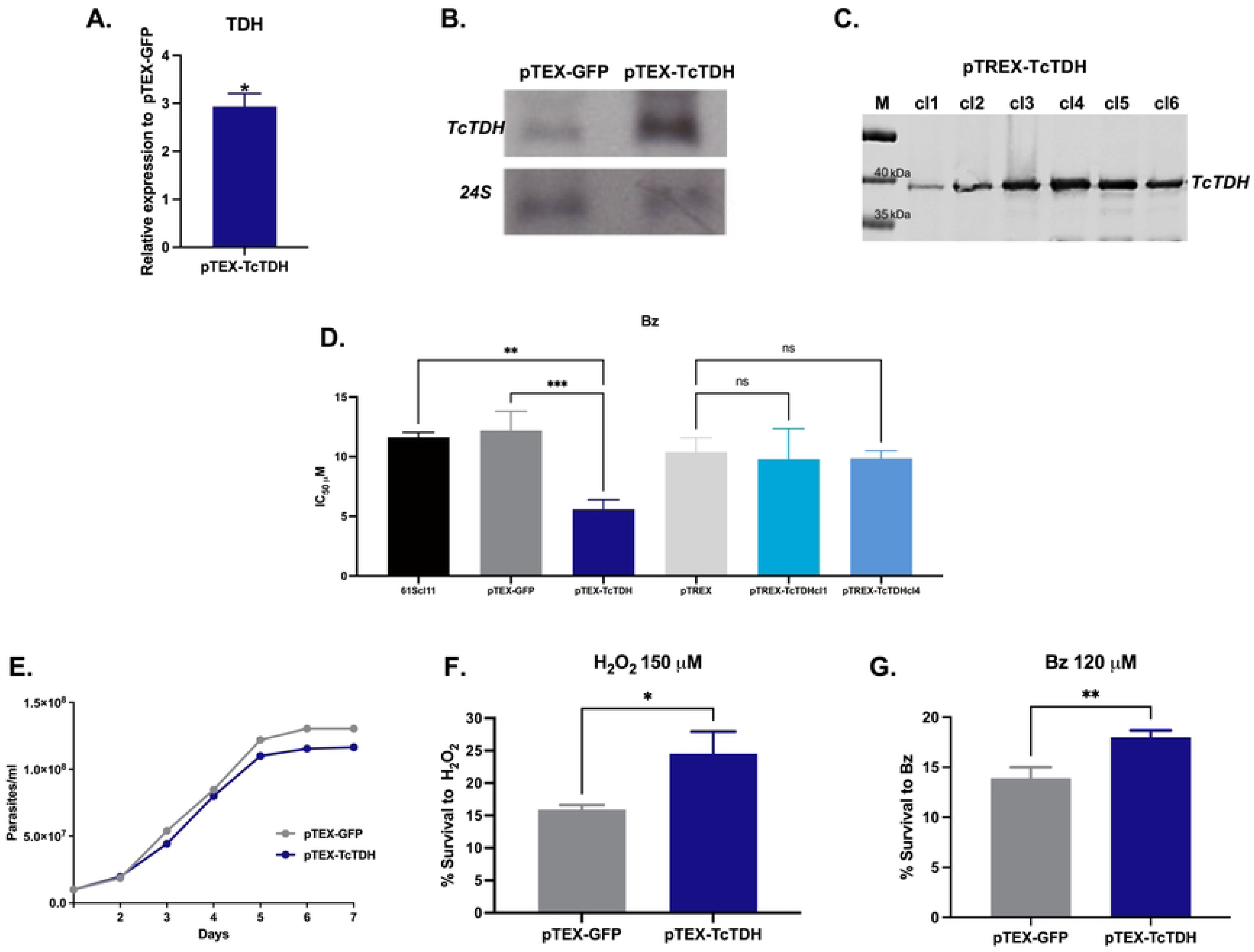
**A.** Relative quantification by RT-qPCR of TcTDH transcript in parasites overexpressing the gene, compared with pTEX-GFP parasites. The mRNA levels were normalized using the expression of the reference gene (HGPRT). The relative quantification results were obtained using the REST2009 program by randomization test with 10,000 permutations (p < 0.05). The graph shows the data in triplicate and two independent experiments. **B.** Northern blot of TDH overexpressing parasites compared with parasites transfected with pTEX-GFP. The ribosomal 24S gene was used as the normalizer. **C.** Western blot from *T. cruzi* clones (cl1-cl6) overexpressing the TcTDH HA-tagged protein in the pTREX vector. M: protein ladder. Anti-HA (1:1000) was used as the primary antibody, and the IRDye 800 (1:15,000) CW Donkey anti-rabbit was secondary. **D.** The IC_50_ to Bz evaluated by alamarBlue for parasites overexpressing the TcTDH gene (pTEX-TcTDH, pTREX-TcTDHcl1 and pTREX-TcTDHcl4), compared with control parasites (61Scl11, parasites transfected with pTEX-GFP or pTREX). Statistical significance was determined in GraphPad Prism 10 using one-way ANOVA with Tukey’s multiple comparison test. **D.** Epimastigotes proliferation curves assessed by counting in Neubauer chamber every 24 h for seven days. **F-G.** Survival curves of parasites overexpressing the TcTDH gene compared to parasites transfected with GFP after de treatment with 120 µM of Bz (F) or 150 µM of H_2_O_2_. Statistical significance was determined in GraphPad Prism 10 using an unpaired t test. ****p < 0.0001; ***p < 0.001; **p < 0.01; *p < 0.05.

**S3 Fig.**
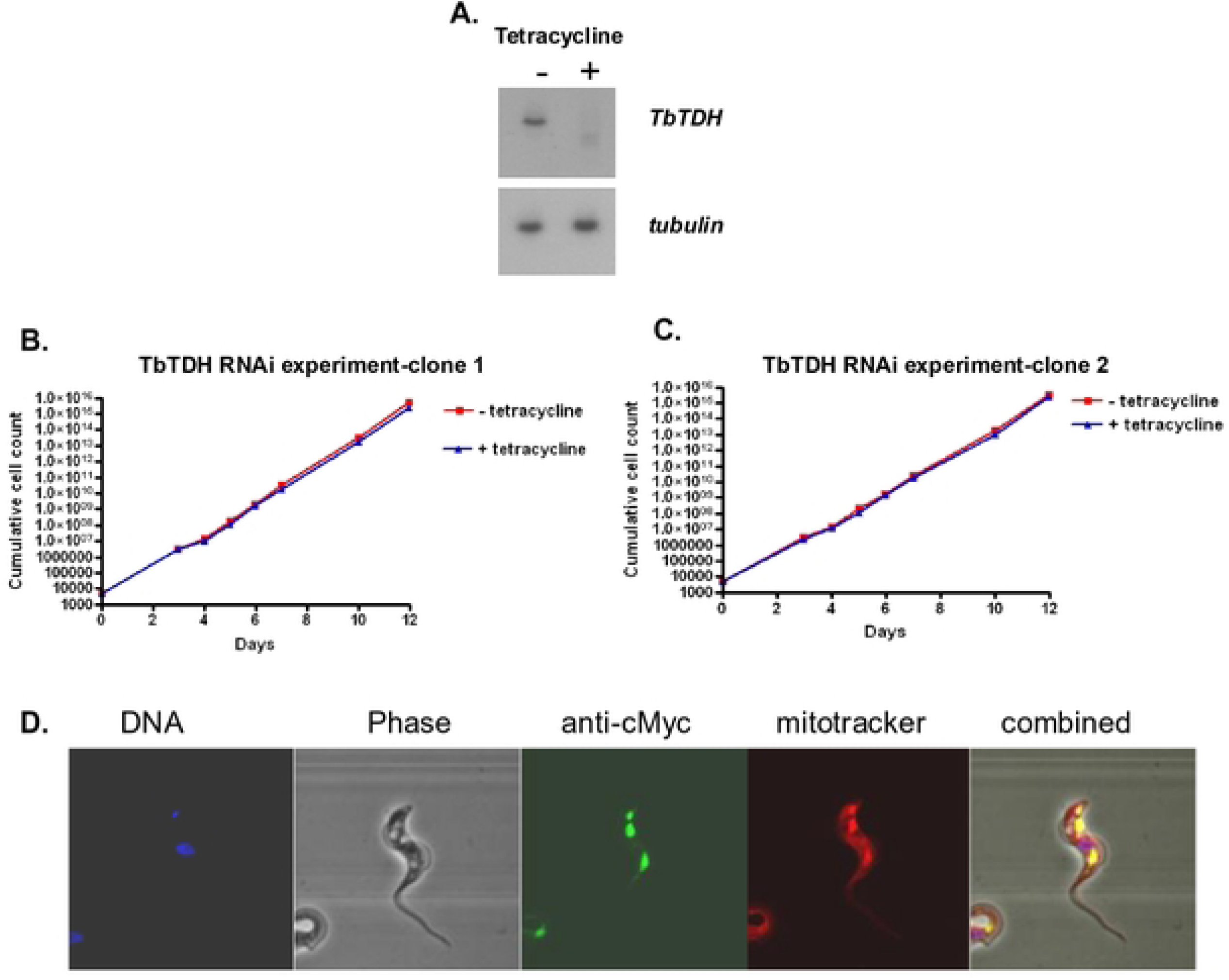
**A.** Northern blot to confirm the knock-down of the TbTDH transcript after 24 hours exposure to tetracycline. The tubulin gene was used as a load control. **B and C**. Growth experiments carried out on two independent TbTDH RNAi lines. There was no significant difference in cumulative growth after 12 days in the presence/absence of tetracycline. **D.** Immunofluorescence microscopy localization using a sequence encoding the 9E10 epitope (derived from human c-Myc) in *T. brucei*. The parasites were incubated with MitoTracker™ (red), and as a primary antibody, anti-cMyc was used and as secondary anti-mouse Alexa 488 (green), with DAPI (blue). The slides were analyzed on the confocal microscope.

---

**S1 Table.** Sequence of primers used in this study.

**S1 File.** Read counts and Differentially regulated genes (DRGs). Read counts obtained from Bz-sensitive and resistant *Trypanosoma cruzi* clones. 61R_cl4: resistant clone; 61S: sensitive clone; LER and WTS: resistant and sensitive clones, obtained by Lima et al. (2023) [20]. The letters a, b, and c correspond to the replicates (Sheet ReadCounts). DGR between Bz-sensitive (61S) vs. Bz-resistant (61R_cl4) *Trypanosoma cruzi* clones (Sheet 61Svs61R) and DGR between Bz-sensitive (WTS) vs. Bz-resistant (LER) *Trypanosoma cruzi* clones (Sheet WTSvsLER). Up- and down-regulated genes are highlighted in blue and red, respectively.

